# scSurv: a deep generative model for single-cell survival analysis

**DOI:** 10.1101/2024.12.10.627659

**Authors:** Chikara Mizukoshi, Yasuhiro Kojima, Shuto Hayashi, Ko Abe, Daisuke Kasugai, Teppei Shimamura

## Abstract

Single-cell omics analysis has unveiled the heterogeneity of various cell types within tumors. However, no methodology currently reveals how this heterogeneity influences cancer patient survival at single-cell resolution. Here, we introduce scSurv, combining a Cox proportional hazards model with a deep generative model of single-cell transcriptome, to estimate individual cellular contributions to clinical outcomes. The accuracy of scSurv was validated using both simulated and real datasets. This method identifies cells associated with favorable or adverse prognoses and extracts genes correlated with their contribution levels. In melanoma, scSurv reproduces known prognostic macrophage classifications and facilitates hazard mapping through spatial transcriptomics in renal cell carcinoma. We also identified genes consistently associated with prognosis across multiple cancers and demonstrated the applicability of this method to infectious diseases. scSurv is a novel framework for quantifying the heterogeneity of individual cellular effects on clinical outcomes.

## INTRODUCTION

Survival analysis is a statistical method widely used across various fields to model the time until a specific event occurs, accounting for the influence of multiple covariates. These models typically consist of two primary components: the baseline hazard function, which represents the underlying hazard when all covariates are absent, and the effect parameters, which quantify how explanatory covariates influence the hazard function. Among them, the Cox proportional hazards model is the most widely used. This semi-parametric model assumes covariates have a multiplicative effect on the hazard function and allows the estimation of effect parameters without specifying the form of the baseline hazard function.

The Cox proportional hazards model has significantly enhanced the analysis of associations between molecular profiles-obtained through next-generation sequencing (NGS) and mass spectrometry-and survival times, enabling the identification of prognostic factors and the development of predictive models ^1–3^. However, traditional approaches based on population averages are limited, particularly when addressing the cellular heterogeneity inherent in diseases and the specific role of various tissues and cell types in pathology. The diversity of cells and intercellular interactions within tumors and the tumor microenvironment influences disease progression and treatment responses ^4–6^, underscoring the need for more precise analyses at the cellular level.

Recent advances in single-cell sequencing technologies have enabled the high-resolution characterization of gene expression patterns and individual cell states, offering novel insights into disease mechanisms and identifying therapeutic targets ^7–9^. Nevertheless, large-scale cohorts integrating single-cell data with clinical information remain limited due to technical and economic constraints. In contrast, bulk RNA sequencing data linked to clinical outcomes are widely available. Despite their lower resolution, these data are valuable for analyzing relationships between molecular profiles and clinical outcomes.

Recent computational advancements have enabled the estimation of cell type proportions from bulk RNA sequencing data ^10–12^. However, deconvolution models based on cell types inherently restrict analyses to this level, failing to capture variations in individual cell states or functions. These limitations hinder our ability to elucidate detailed biological mechanisms, including subtle shifts in cell states or associations between specific subtypes and survival outcomes. Consequently, critical biological insights may be overlooked, particularly in complex diseases like cancer, which exhibit pronounced intratumoral heterogeneity.

We addressed these limitations by developing scSurv, a novel method for single-cell survival analysis that extends the Cox proportional hazards model to incorporate cellular heterogeneity. Using single-cell RNA sequencing data as a reference, scSurv decon-volves bulk RNA-seq data to infer single-cell proportions and quantifies their individual contributions to patient outcomes. scSurv systematically identifies cells contributing to disease risk and their specific gene signatures by analyzing the association between survival time and cell abundance.

Simulations demonstrated that scSurv accurately predicts patient prognosis using single-cell proportions derived from deconvolution. Applying this method to The Cancer Genome Atlas (TCGA) data, we found that it could predict survival time of patients excluded from training across multiple cancers. scSurv identified specific cells influencing patient prognosis and the genes associated with these outcomes in melanoma. It also successfully reproduced known macrophage classifications affecting patient prognosis. Spatial transcriptomic analysis of renal cell carcinoma enabled tissue-wide hazard mapping and identified distinct prognosis-associated spatial regions. Furthermore, we identified genes contributing to prognosis across multiple cancers. Finally, applying this method to infectious diseases demonstrated its utility beyond cancer and survival outcomes. This novel approach elucidates the contributions of individual cells to clinical outcomes and offers new perspectives in clinical analyses.

## RESULTS

### Concept of scSurv

We developed a novel method called scSurv, a deep generative model for single-cell survival analysis, to facilitate survival analysis at the single-cell level and to uncover biological mechanisms in diseases (Figure 1). scSurv involves the following steps: 1. This method uses single-cell RNA sequencing (scRNA-seq) data as a reference, following our previous framework ^13^. Bulk RNA sequencing (bulk RNA-seq) data are deconvoluted at the cellular level using latent cell states obtained from a variational autoencoder (VAE). 2. scSurv estimates the hazard function using a Cox proportional hazards model, extended by combining the estimated proportion of each single cell within the bulk samples and the regression coefficients obtained from the latent cell state. These regression coefficients are interpreted as the contributions of individual cells to clinical outcomes. This model enables the evaluation of hazard contributions at the single-cell level and enhances their consistency among cells with similar cell states.

**Figure 1:**
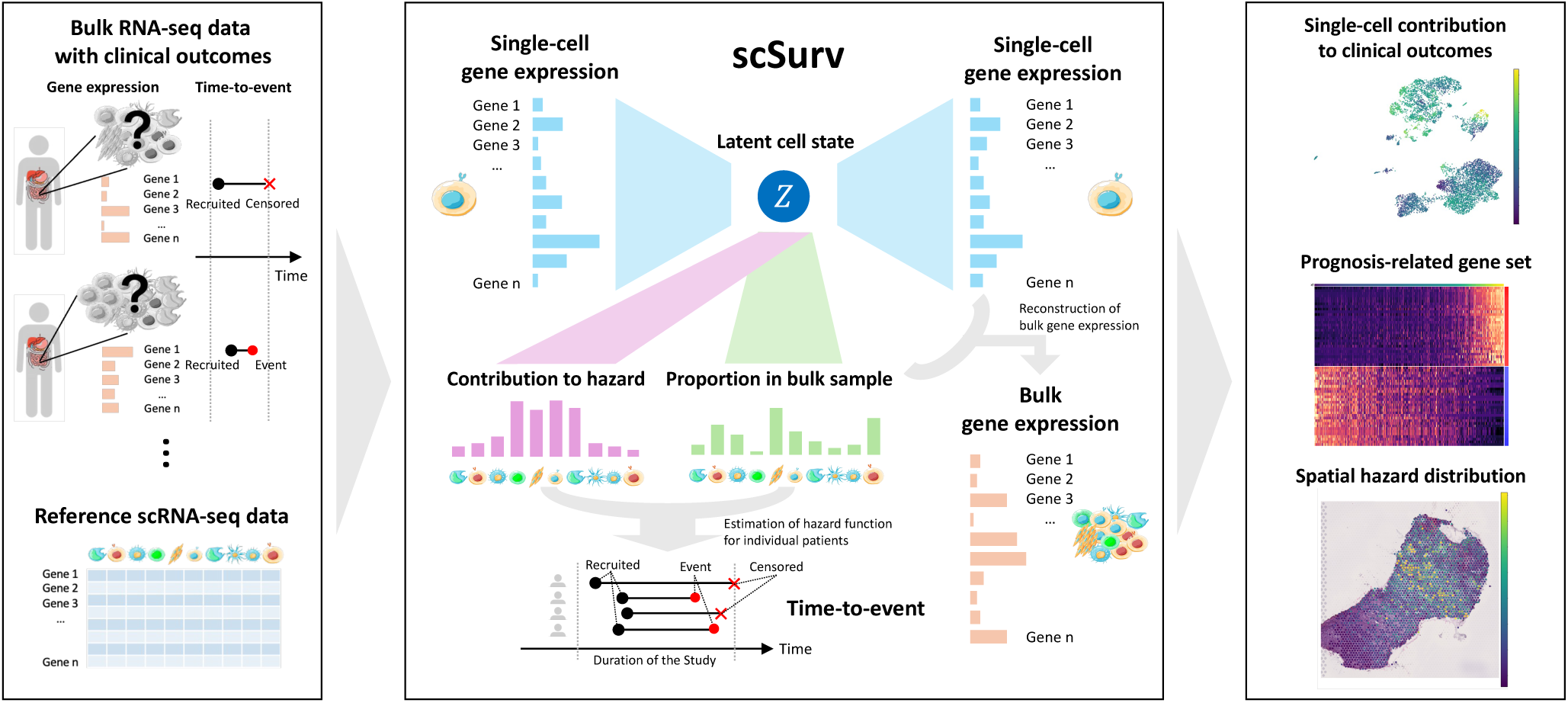
Overview of scSurv. Overview of scSurv, a deep generative model for single-cell survival analysis. This method deconvolutes bulk RNA-seq data into each single cell using VAE, followed by survival analysis with an extended Cox proportional hazards model. The framework enables single-cell level prognostic analysis, identification of outcome-associated genes, and spatial hazard mapping.

The trained model offers three main analyses: 1. Quantification of individual cells’ contributions to clinical outcomes 2. Identification of prognosis-associated gene sets 3. Mapping of spatial hazard distributions using spatial transcriptome data

Through these applications, scSurv provides a comprehensive and interpretable framework that reveals heterogeneity in the clinical significance of cells. This framework enables the identification of novel cell populations and genes involved in the prognosis. This method is available as an open-source Python package on GitHub (https://github.com/3254c/scSurv).

### Validation using simulated datasets

We validated the performance of scSurv using simulated datasets. Whereas the existing bulk RNA-seq deconvolution methods only perform cluster-level resolution, scSurv performs deconvolution at the single-cell level. Through simulations, we assessed the accuracy of scSurv’s deconvolution and the precision of estimating the contributions to the hazard function based on the inferred cell proportions. Our results showed that under realistic conditions, single-cell level deconvolution achieves higher accuracy and is more effective for hazard regression than cluster-level methods.

We first clustered scRNA-seq data and assigned cluster labels to generate the simulated data. We then split the data into two subsets: one used as a reference and the other used to create pseudo-bulk samples. We randomly determined the proportion of each cell cluster within the bulk samples and generated pseudo-bulk RNA-seq data by aggregating their expression. We assigned regression coefficients in the hazard function to each cluster and set the survival times for each pseudo-bulk sample based on the Cox proportional hazards model.

We evaluated scSurv’s performance in deconvoluting bulk data and compared it with existing methods such as CIBERSORTx ^10^, MuSiC ^11^, and BayesPrism ^12^, which perform cluster-level deconvolution of bulk RNA-seq using scRNA-seq data as a reference. However, these methods do not achieve single-cell resolution. In realistic settings, the boundaries of meaningful cell populations may not align with the provided cluster labels. Therefore, we evaluated two scenarios: one in which we provided the same cluster labels used to generate the pseudo-bulk data for these methods and the other in which we provided labels with different boundaries.

The cell-type proportions estimated by scSurv were positively correlated with the ground-truth values in the simulation(Figure 2A). When the existing methods were given the same cluster labels used to generate pseudo-bulk RNA-seq data, their estimated cluster-level proportions were more accurate than those of scSurv. However, when different labels were provided, scSurv’s decon-volution accuracy was higher than the existing methods. This finding indicates that cluster-level deconvolution methods are limited by performance variability depending on the cluster labels provided and the clustering process. In contrast, scSurv’s single-cell level deconvolution avoids the bias introduced by clustering and produces consistent results.

**Figure 2:**
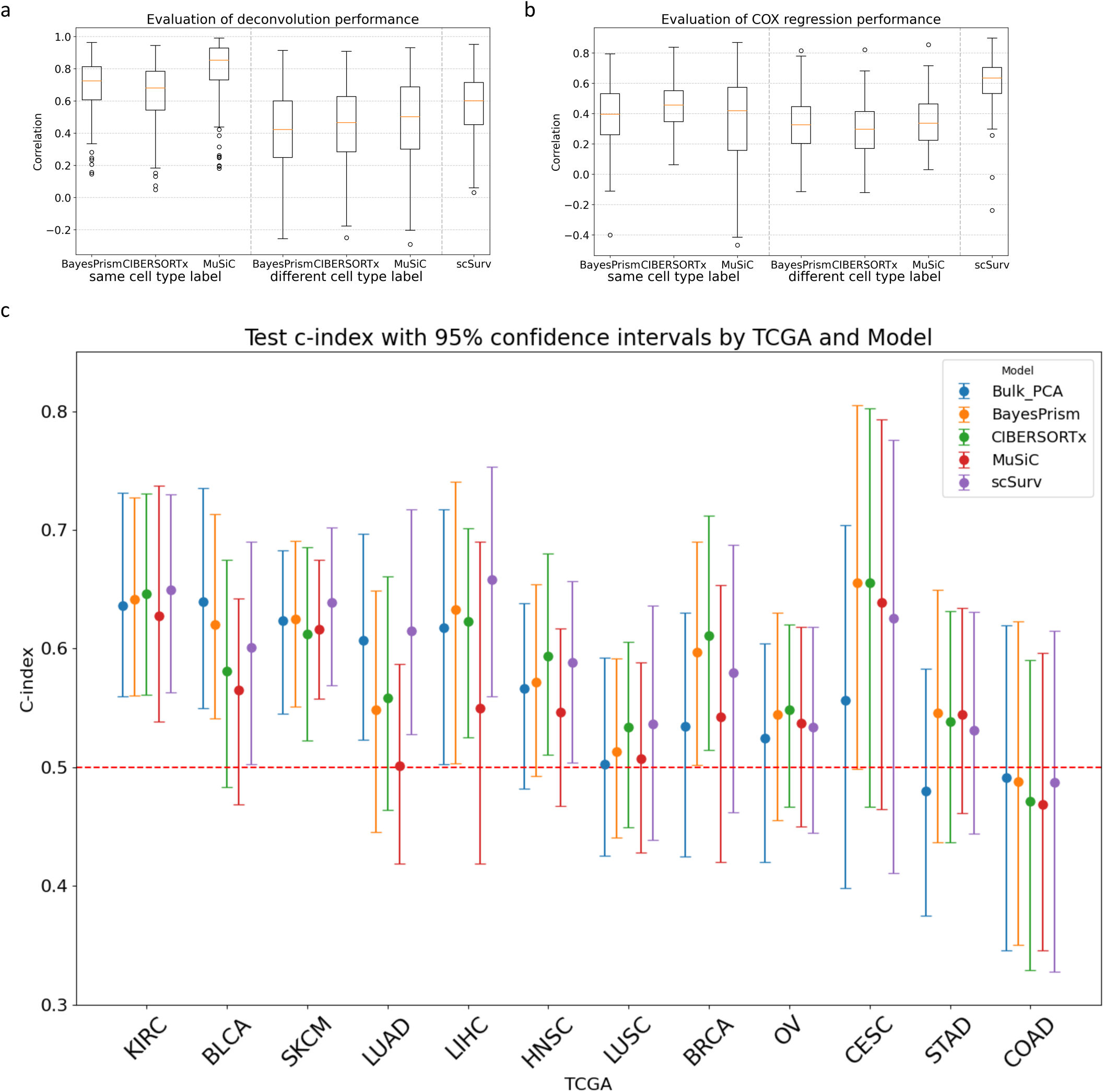
Evaluation of scSurv. (a) Deconvolution accuracy of scSurv, BayesPrism, CIBERSORTx, and MuSiC using 300 simulated pseudo-bulk samples. Correlation between estimated proportions and ground truth values was calculated. BayesPrism, CIBERSORTx, and MuSiC were evaluated under two conditions: using same or different cluster labels between pseudo-bulk generation and deconvolution. (b) Accuracy of regression coefficient estimation using simulated data across methods. Correlation between estimated and ground truth regression coefficients was calculated across 100 datasets, which were generated using the same 300 pseudo-bulk samples with different survival time settings by varying coefficient values. For BayesPrism, CIBERSORTx, and MuSiC, coefficients were estimated using the lifelines library based on deconvoluted cell type proportions under two conditions. (c) The performance of scSurv, the combination of the existing methods for bulk deconvolution with the Cox proportional hazards model, and the combination of bulk PCA with the hazard model were evaluated across 12 TCGA cancer types using the c-index. 95% confidence intervals were calculated from 100 iterations with different train/validation/test splits. scSurv achieved a test c-index >0.5 in KIRC, BLCA, SKCM, LUAD, LIHC and HNSC. Lower performance in certain cancers suggests insufficient prognostic information in bulkRNA-seq data.

Additionally, when estimating the contribution of each cluster to the hazard function using the inferred cell proportions, scSurv’s estimates were positively correlated with the ground-truth values (Figure 2B). In both cases, where the same cluster labels were used as those in generating the pseudo-bulk RNA-seq and where different labels were used, scSurv’s estimation accuracy for contributions was higher than that of the existing methods. We found that cluster-level deconvolution methods produce results that vary depending on the clusters provided and fail to accurately estimate contributions when the cluster labels used in the hazard estimation differed from those used to determine hazard values in the simulation. These results validate the accuracy of scSurv’s estimations and support the advantage of performing single-cell level estimation of the contribution to the hazard function.

### Evaluating generalization performance using TCGA datasets

We applied scSurv to real datasets from TCGA. We selected 12 cancer types that had bulk RNA-seq data from over 300 patients to ensure learning stability. In the training data, scSurv successfully predicted the hazard function based on cell proportions (Figure S1D). In the test data, scSurv demonstrated generalization performance by achieving a 95% confidence interval for the concordance index above 0.5 in multiple cancers (Figure 2C). However, in some cancer types, the 95% confidence interval for estimation accuracy crossed 0.5, indicating difficulty in prediction. In these cancers, the confidence intervals of the c-index for the hazard estimation also crossed 0.5, even when we directly used the bulk RNA-seq data for hazard estimation, suggesting that bulk RNA-seq data may lack sufficient information to accurately predict the hazard function. This observation is consistent with previous studies that reported challenges in predicting prognosis using TCGA bulk RNA-seq data for certain cancer types ^14,15^. Moreover, for the cancers with high estimation accuracy using bulk RNA-seq data (the lower bound of 95% CI > 0.5), scSurv’s estimation accuracy outperformed the existing methods, except for BLCA. For the cancers with low prediction accuracy (the lower bound of 95% CI < 0.5), we found that the 95% CI of the c-index crossed 0.5 for most of the methods, suggesting that meaningful estimation of the hazard and their comparison were difficult in these cancers. From these findings, scSurv achieved accurate estimations across several cancer types, indicating that single-cell wise estimation of the hazard contribution by scSurv leads to more accurate prognosis prediction from bulk transcriptome data.

### Identifying cells and genes associated with melanoma prognosis

We applied scSurv to a melanoma cohort (TCGA-SKCM). Melanoma is the most lethal type of skin cancers worldwide. ^16^ Our analysis revealed that specific populations of cancer cells, fibroblasts, endothelial cells, and macrophages adversely affected survival outcomes (Figure 3A, Figure 3B). We permuted the estimated contributions across cells within each cell cluster and quantified the decrement in the c-index to determine which cell clusters are important for prognostic prediction (Figure 3C). Here, we found that cancer cells and macrophages exhibited a larger decrement, indicating that the heterogeneity of hazard contributions within these cell types had larger effects on the prognosis than the other cell types. Although intratumoral and intertumoral heterogeneity of cancer cells are documented factors influencing patient outcomes ^17^, recent findings additionally highlight the critical role of tumor-associated macrophages in shaping the tumor microenvironment and affecting patient prognosis ^18^. Accordingly, our analysis focused on macrophages.

**Figure 3:**
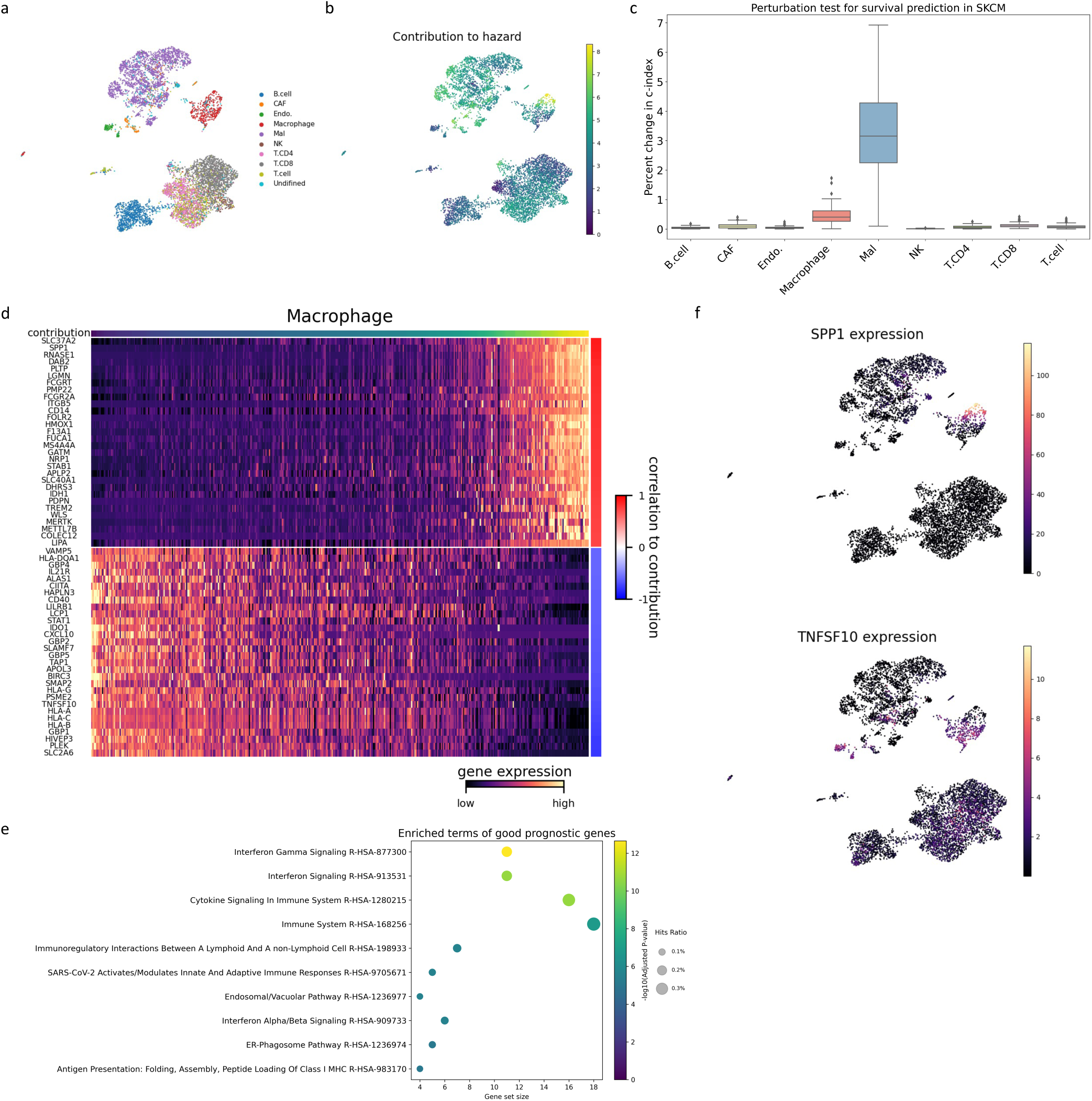
scSurv reveals prognostic cells and genes in melanoma. (a) UMAP visualization of cell type annotations in melanoma dataset. (b) Single-cell contributions estimated by scSurv. Single-cell contributions are regression coefficients in the hazard function, which are estimated using single-cell proportions; both the contributions and proportions are derived from the latent states of a VAE. Higher values indicate adverse prognostic impact. (c) Permutation test identifying prognostically important clusters. Estimated hazard contributions were permuted across cells within each cluster, and the resulting decrement in the c-index was quantified to assess the impact of each cluster on hazard estimation. (d) Heatmap showing expression of genes correlated with estimated contributions. The expression reconstructed from the latent variables was used. Top 30 positively and negatively correlated genes are displayed. The estimated single-cell contributions are shown in the upper panel. (e) Dot plot showing gene set enrichment analysis results using the top 30 negatively correlated genes with contributions. Dot size represents the proportion of enriched genes, the color indicates statistical significance, and the x-axis shows the number of genes in each pathway. (f) UMAP visualization of reconstructed SPP1 and TNFSF10 expression, showing concordance with their prognostic contributions.

We isolated macrophages and identified genes whose expression correlated with scSurv estimated contributions (Figure 3D). Gene set enrichment analysis revealed that interferon gamma signaling pathways were associated with favorable prognosis (Figure 3E). This finding is consistent with those of previous studies. ^19^ Additionally, the SPP1 gene characterizes macrophage subsets that contribute to tumor promotion. ^20^ In contrast, the TNFSF10 gene encodes TNF-related apoptosis-inducing ligand (TRAIL), which promotes antitumor activity by inducing tumor cell apoptosis ^21,22^. In scSurv’s estimations, SPP1 expression positively correlated with adverse prognosis, whereas TNFSF10 expression correlated negatively (Figure 3F). These results indicate that scSurv can classify macrophages into prognostic subsets, aligning with previous studies.

Our results confirm scSurv effectiveness in identifying prognostically relevant cells and genes. The consistency of these findings with existing knowledge highlights scSurv’s utility in biological analyses.

### Integration with spatial transcriptomics data in renal cell carcinoma

We expanded the scSurv analysis by integrating spatial transcriptomics data. First, we conducted the standard scSurv estimation to assign contributions to each cell using a renal cell carcinoma cohort (TCGA-KIRC) and scRNA-seq data ^23^ (Figure 4A, Figure 4B). Next, we estimated the cellular composition for each spot in the spatial transcriptome. We assigned a spatial hazard score to each spot based on single-cell proportions and individual cell contributions (Figure 4C). Regions with high spatial hazards were distributed between areas containing cancer cells and those containing normal cells(Figure S2A).

**Figure 4:**
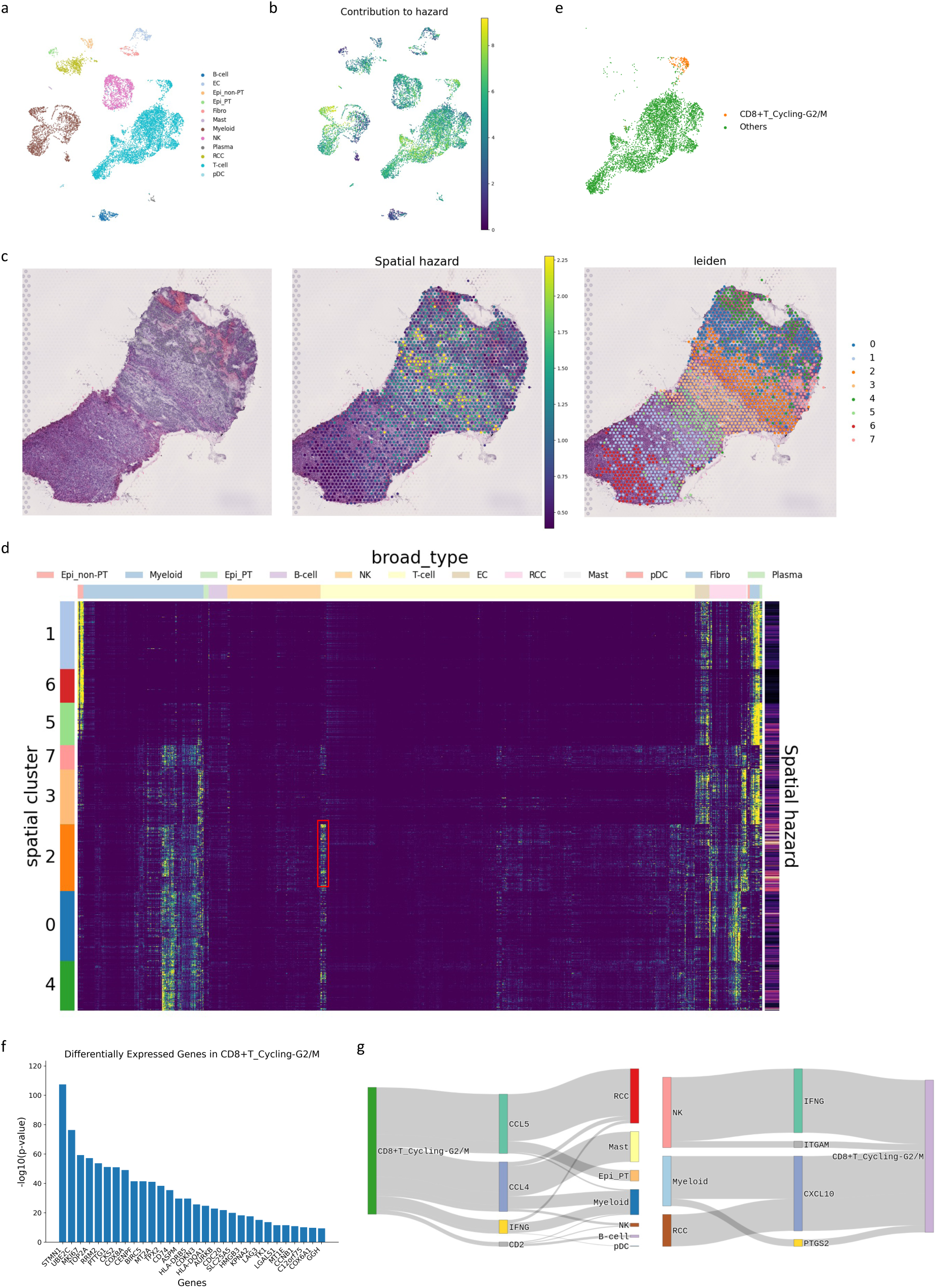
Integrated analysis using spatial transcriptomics in renal cell carcinoma. (a) UMAP visualization of cell type annotations in renal cell carcinoma dataset. (b) Single-cell contributions estimated by scSurv. Higher values indicate adverse prognostic impact. (c) Spatial visualization showing H&E image, mapped spatial hazards, and spot-level clustering. The minimum or maximum hazard value of the top 2.5% was forced to the 2.5% and 97.5% quantile values to prevent extreme values from affecting the visualization. (d) Heatmap of single-cell contributions adjusted for cell proportions. Cluster 2 showed the highest spatial hazard, with specific T cell populations. (e) UMAP visualization of cluster 2 specific T cell populations. (f) Differentially expressed genes in identified T cell clusters. (g) Sankey plot showing cell cell communications between the identified T cell cluster and other cell types predicted by NicheNet.

We clustered the spots based on their expressions to obtain spatial clusters, calculated the average hazard for each spatial cluster, and focused on clusters with particularly high hazards (Figure 4D). Notably, we found that proliferative CD8-positive T cells specifically influenced the hazard function within this cluster (Figure 4E, Figure S2B, Figure S2C, Figure S2D). This population of T cells is characterized by the expression of the Ki-67 gene (Figure 4F) and is associated with poor prognosis in renal cell carcinoma ^24^. Analysis of intercellular communication networks involving spatially co-localized cells in this population identified several prognostically relevant genes, including CCL4, CCL5, IFNG, ITGAM, and CXCL10^25–29^ (Figure 4G). Integrating scSurv with spatial transcriptomics allowed us to perform hazard mapping across tissue sections, identify specific areas linked to patient prognosis, and examine the interactions between key cell populations.

### Multiple cancer analysis

We performed a scSurv analysis of multiple cancer types. We selected six cancers where the lower bound of the 95% CI exceeded 0.5 in test patients and analyzed the cell populations and genes consistently linked to survival outcomes across these cancers. We first conducted permutation tests on the contribution scores of common cell types across the six cancers, including endothelial cells, fibroblasts, T cells, B cells, and myeloid cells, to determine their relative importance in the hazard function prediction (Figure 5A). Among the analyzed cell types, myeloid cells had the strongest influence on prediction accuracy. We extracted myeloid cell populations from each cancer type and examined their correlations with the contributions of genes commonly expressed across all cancer types (Figure 5B). We performed gene set enrichment analysis on the top and bottom genes ranked by mean correlation to elucidate the biological pathways consistently associated with prognosis in myeloid cells across different cancers. Genes associated with a good prognosis were enriched in antigen presentation-related terms (Figure 5C), consistent with the known role of antigen presentation in antitumor immunity. ^30^

**Figure 5:**
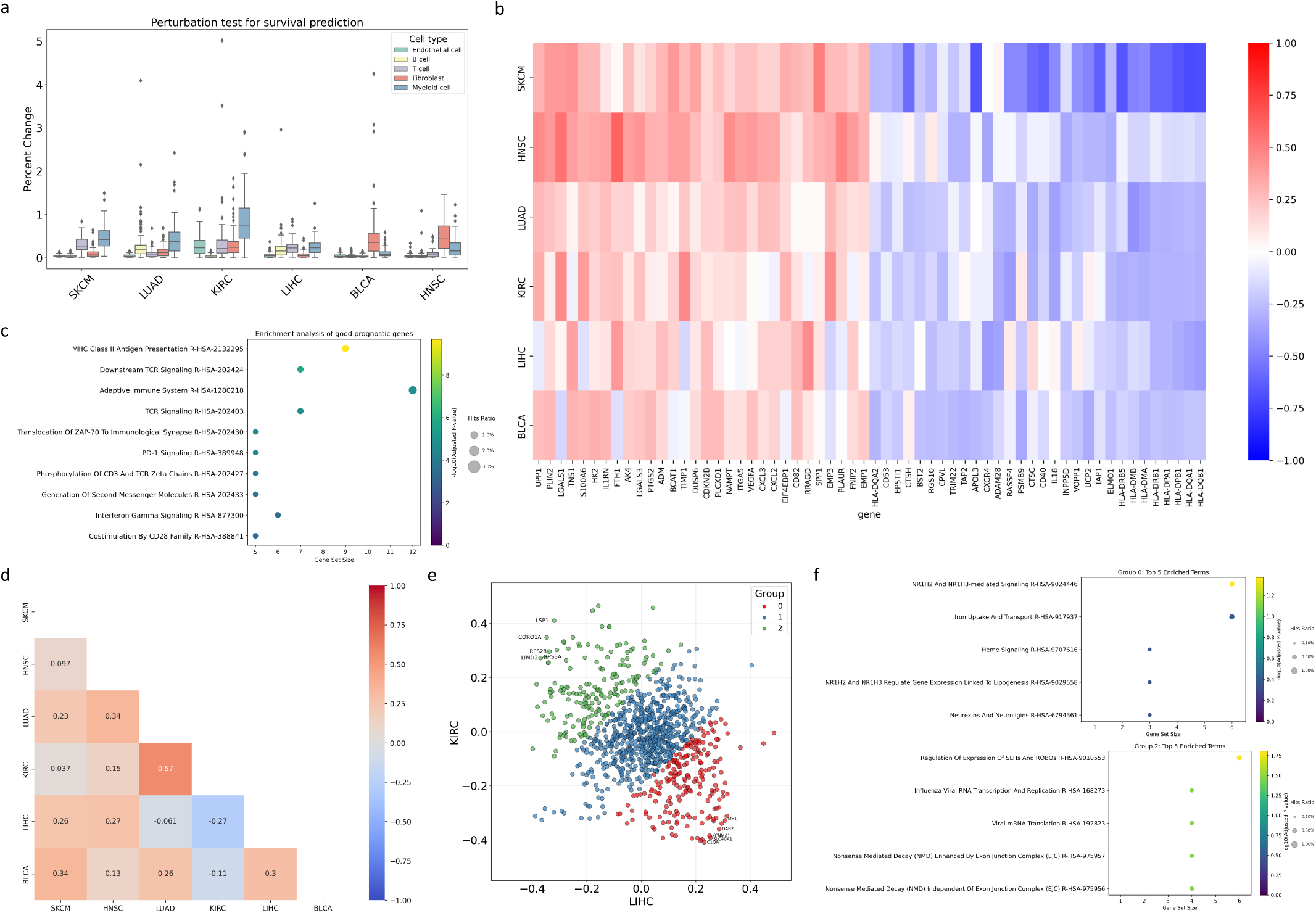
Pan-cancer analysis of stromal cells. (a) Permutation test identifying common prognostic cell types across six cancer types. (b) Heatmap showing correlations between myeloid cell gene expression and contributions across six cancer types. The heatmap illustrates the top 30 genes with the highest average correlation and the bottom 30 genes with the lowest average correlation across the cancer types. (c) Dot plot showing gene set enrichment analysis results using the top 30 negatively correlated genes with contributions. Dot size represents the proportion of enriched genes, the color indicates statistical significance, and the x-axis shows the number of genes in each pathway. (d) Cancer similarity based on gene correlations. (e) Scatter plot of correlations between gene expression and contributions in liver and kidney cancer myeloid cells. The genes were clustered into three groups by applying a Gaussian mixture model with three components to their projections onto the *y* = *−x* line. (f) Gene set enrichment analysis results for differentially prognostic genes in liver and kidney cancer myeloid cells.

Next, we calculated the similarities between the cancer types based on their correlations (Figure 5D). Notably, myeloid cells from hepatocellular carcinoma and renal cell carcinoma showed opposite correlation patterns. Analysis of differentially correlated genes (Figure 5E) revealed enrichment of the NR1H2/NR1H3 and Slit/Robo signaling pathways (Figure 5F). These findings suggest that these pathways differentially affect survival outcomes in hepatocellular and renal carcinomas through mechanisms mediated by myeloid cells.

### Application to other clinical outcomes in a COVID-19 Cohort

We applied scSurv to hospitalized COVID-19 patients to demonstrate the applicability of our method to acute immune responses beyond cancer and to clinical outcomes beyond survival. Bulk RNA-sequencing data of peripheral blood mononuclear cells (PBMCs) and the corresponding clinical information were obtained from the IMPACC cohort. ^31^ Single-cell RNA sequencing data from PBMCs of 12 patients in Yoshida et al. ^32^ were used as a reference dataset (Figure 6A). We applied scSurv to two distinct clinical outcomes: survival and discharge. scSurv provided robust predictions for both endpoints (Figure 6B). Here, we found that monocytes exhibited a large contribution to survival hazard(Figure 6C). Cellular contributions to survival hazard exhibited an inverse correlation with discharge hazard (Figure 6D), reflecting the clinical relationship between shortened survival and extended hospitalization in severe disease. Permutation testing identified monocytes as the key cellular population in outcome prediction (Figure 6E). These findings, including the large contribution of monocytes to the survival hazard and their importance identified through permutation testing, align with previous studies demonstrating the importance of monocytes in COVID-19 severity ^33,34^. Based on these observations, we focused our subsequent analyses on this subset.

**Figure 6:**
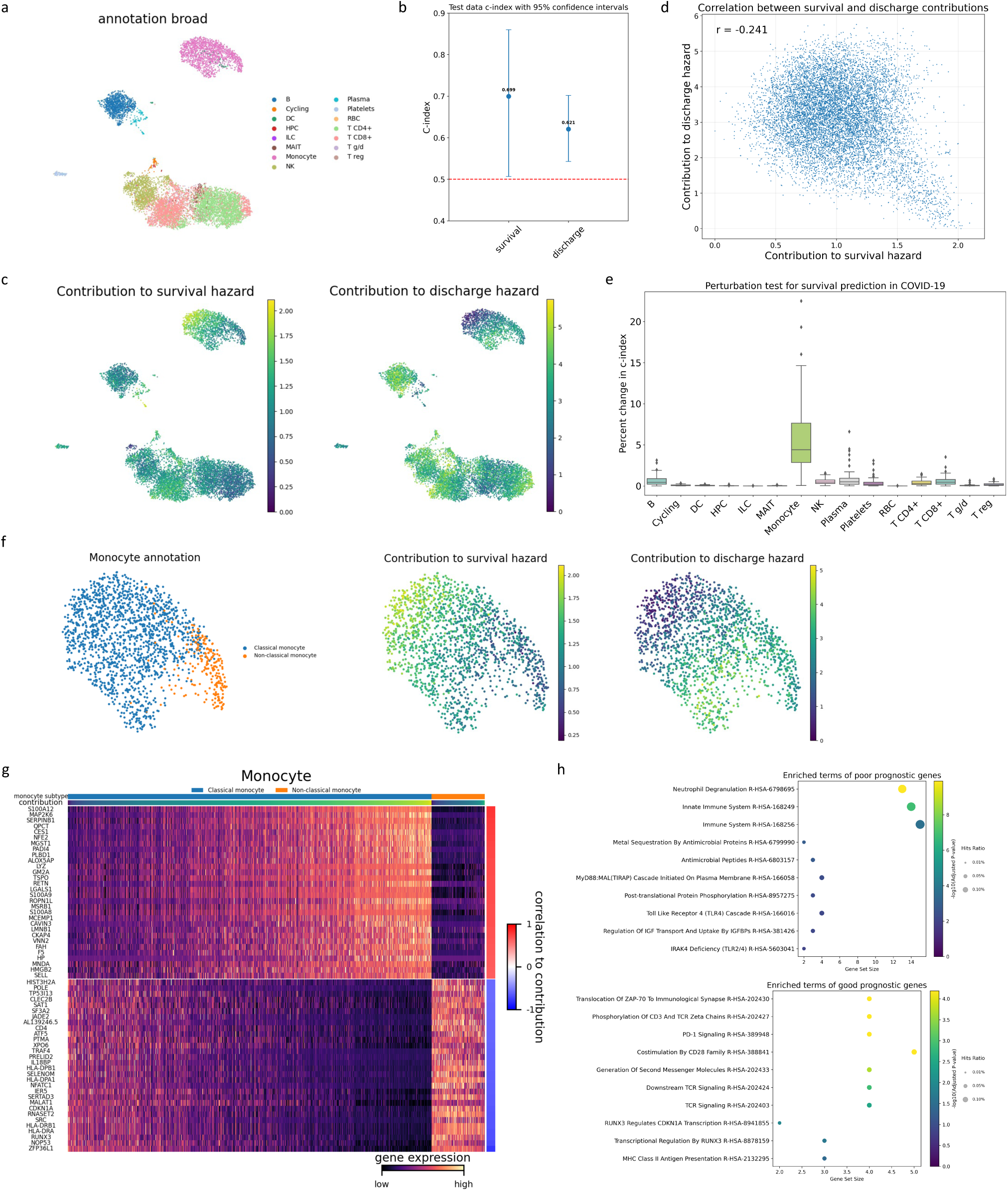
COVID-19 PBMC analysis. (a) UMAP visualization of cell type annotations in COVID-19 dataset. (b) 95% confidence intervals of scSurv c-indices for survival and discharge in test patients. (c) UMAP visualization of single-cell contributions to survival and discharge hazard. Higher values indicate adverse prognostic impact. (d) Scatter plot of single-cell contributions for both outcomes. Cellular contributions to survival hazard exhibited an inverse correlation with discharge hazard. (e) Permutation test identifying prognostically important clusters in COVID-19. (f) UMAP visualization of monocyte annotations and their hazard contributions to both outcomes. Classical monocytes show high contributions to survival hazard but low contributions to discharge hazard. (g) Heatmap showing reconstructed expression of genes correlated with estimated contributions to survival hazard. Top 30 positively and negatively correlated genes are displayed. The estimated single-cell contributions are shown in the upper panel. (h) Dot plot showing gene set enrichment analysis results using the top 30 positively and negatively correlated genes with contributions to survival hazard. Dot size represents the proportion of enriched genes, the color indicates statistical significance, and the x-axis shows the number of genes in each pathway.

The distribution of contributions between classical and non-classical monocyte subtypes aligned with previous findings on the role of classical monocytes in COVID-19 severity ^34^(Figure 6F). The top 30 genes positively correlated with monocyte contributions included S100A12, S100A9, and S100A8, while the top 30 genes negatively correlated with monocyte contributions included HLA-DRA, HLA-DRB1, HLA-DPA1, and HLA-DPB1(Figure 6G). High expression of S100A proteins and low expression of HLA-DR proteins in monocytes have been identified as hallmarks of progressive disease. ^35^ These gene expression patterns were previously linked to disease severity in an analysis of scRNA-seq data from 66 PBMC samples. ^36^ While collecting scRNA-seq data from a large number of individuals is cost-prohibitive, scSurv enables single-cell level analyses by integrating bulk RNA-seq data with a smaller scRNA-seq reference dataset. Our method provides a cost-effective solution for high-resolution profiling. Gene set enrichment analysis further revealed that genes positively correlated with monocyte contributions were enriched in neutrophil degranulation pathway and innate immune system pathway, aligning with reports linking activation of innate immune cells to poor COVID-19 prognosis(Figure 6H). ^37,38^ Conversely, genes negatively correlated with monocyte contributions were enriched in pathways related to TCR signaling and antigen presentation, supporting findings that impaired adaptive immune responses and antigen presentation are associated with disease severity. ^39,40^ These findings highlight scSurv’s broader applicability beyond cancer and its utility in analyzing time-to-event data beyond survival time.

## DISCUSSION

We present the first methodology to quantify individual cells’ contributions to clinical outcomes. This method allows the investigation of cells linked to patient outcomes using existing cohorts such as TCGA, enabling novel insights into cellular heterogeneity at unprecedented resolution. scSurv does not rely on cell clustering, and its estimations are unaffected by cell labels, providing robust and unbiased results. Additionally, by utilizing the framework of conditional variational autoencoders for batch effect removal, we can effectively integrate scRNA-seq data from multiple patients to use as a reference. scSurv extends the Cox proportional hazards model by incorporating both the estimated proportions of single cells in bulk RNA-seq samples and the contributions of individual cells to clinical outcomes, both derived from latent cell states obtained through a VAE. By leveraging these latent variables, this extended model enables precise survival analysis at the single-cell level and effectively scales to analyze contributions across large numbers of cells.

However, we recognize that scSurv has certain limitations. Similar to existing deconvolution methods, our method cannot assign proportions or contributions to cells that are not included in the reference. Therefore, it is essential to select reference datasets that comprehensively represent the cell populations found in bulk RNA-seq data. Additionally, the method requires a sufficient number of patients (over 300) and cannot be applied to cancers with rare death events. Furthermore, in certain cancer types, bulk RNA-seq data may contain insufficient predictive information, making accurate hazard function estimation difficult. Addressing these constraints remains a priority for future methodological development.

Despite these limitations, scSurv is a novel method to quantitatively evaluate individual cells’ effects on clinical outcomes, enable identification of clinically relevant cell populations and genes, and generate new insights through integration with spatial transcriptomics. This method introduces an innovative concept of cellular heterogeneity based on contributions to clinical outcomes.

In summary, scSurv represents a significant advancement in single-cell analysis methodology, bridging the gap between cellular heterogeneity and clinical outcomes. We anticipate that scSurv will become an essential tool for researchers investigating the cellular basis of disease progression and treatment response, ultimately contributing to the development of more effective, personalized therapeutic strategies.

## METHODS

### scSurv framework

The scSurv framework consists of three main steps: learning latent states through VAE, deconvolution of bulk data, and estimation of regression coefficients in hazard functions for each cell using the extended Cox proportional hazards model. The first two steps are similar to those in our previous study, DeepCOLOR ^13^. scSurv employs a conditional VAE framework ^41^ similar to scVI ^42^ to handle scRNA-seq data collected from multiple patients as input. The VAE compresses raw gene expression into low-dimensional latent cell representations, which can be treated as summaries of essential cellular information. scSurv utilizes the loss function of the Cox proportional hazards model to learn the contribution to the hazard function using the proportion of each single cell in each bulk sample as covariates. The proportions of the cells and their contributions to the hazard function learned in this process depends on the latent states of the cells. Consequently, cells with similar latent states are estimated to have similar contributions. This approach enables the model to effectively learn the contributions of 10,000 cells.

### Derivation of the stochastic latent representation of the single-cell transcriptome

We define a probabilistic model for raw counts of single-cell transcriptomes. Let **z***_c_* ∈ ℝ*^M^* represent the low-dimensional latent cell state, where *M* is the dimension of the latent space and the subscript *c* denotes each cell. We assume a Gaussian prior distribution for **z***_c_*:

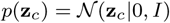

Let *G* be the number of genes and **x***_c_* ∈ ℝ*^G^* be the raw counts of the single-cell transcriptome. We assume that given **z***_c_*, the counts follow a Poisson distribution:

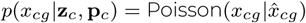

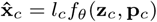

Here, *l_c_* ∈ ℝ is the mean of all genes for each cell, **p***_c_* ∈ [0, 1]*^P^* represents the batch information (patient information in this context), *P* is the total number of batches, and *p_c,k_* = 1 indicates that cell *c* is collected from patient *k*. The function *f_θ_*(**z***_c_,* **p***_c_*) ∈ ℝ*^G^* is the decoding neural network of the latent cell state.

To obtain the latent cell state **z***_c_*, we use a VAE framework to represent the posterior distribution *q*(**z***_c_*|**x***_c_,* **p***_c_*). We assume that the posterior distribution follows a Gaussian distribution:

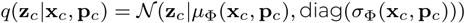

where *µ*_Φ_(**x***_c_*) and *σ*_Φ_(**x***_c_*) denote the encoding neural networks.

We maximize the following Evidence Lower Bound to optimize the parameters of the generative model and variational distribution:

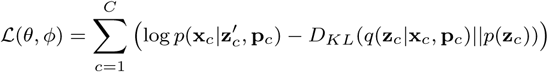

where 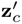 is obtained from *q*(**z***_c_*|**x***_c_,* **p***_c_*) using the reparameterization trick.

### Probabilistic model of bulk transcriptome data and spatial transcriptome data

We model the expression **e***_b_* ∈ ℝ*^G^* of bulk *b* by following a negative binomial distribution:

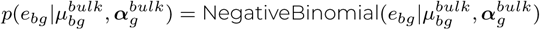

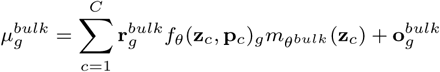

where 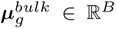 is the mean parameter, *B* is the number of bulk samples, ***α****^bulk^* ∈ ℝ*^G^* is the dispersion parameter, **r***^bulk^* ∈ ℝ*^G^* represents the capture rate of each gene in bulk RNA-seq compared to scRNA-seq, and **o***^bulk^* ∈ ℝ*^G^* is the shift parameter for each gene. *m_θ^bulk^_* (**z***_c_*) ∈ ℝ*^B^* is a neural network that outputs the proportion of each cell in each bulk sample given the latent cell state as the input, satisfying:

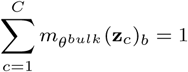

We optimize the parameters *θ^bulk^*, **r***^bulk^*, **o***^bulk^*, *α^bulk^* by maximizing the following log-likelihood:

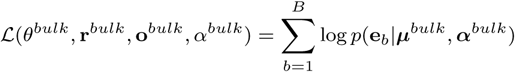

Similarly, we model the expression **e***_s_* ∈ ℝ*^G^* of Visium spot *s* following a negative binomial distribution:

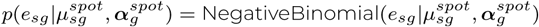

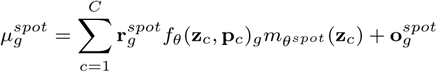

where the parameters are defined analogously to the bulk model, with *m_θ^spot^_* (**z***_c_*) ∈ ℝ*^S^*, where *S* denotes the number of spots, satisfying:

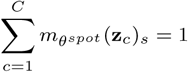

We optimize the parameters *θ^spot^*, **r***^spot^*, **o***^spot^*, *α^spot^* by maximizing the following log-likelihood:

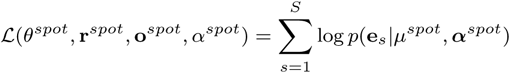

### Single-cell cox proportional hazards model

We define the hazard function for each bulk sample as:

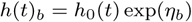

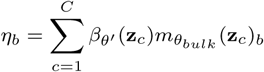

where *β_θ′_* (**z***_c_*) ∈ ℝ*^c^* is a neural network that outputs regression coefficients given latent cell states as input, and *h*_0_(*t*) is the baseline hazard function, which remains unspecified.

Following the Cox proportional hazards model, we optimize the parameters *θ^′^* of *β_θ′_* (**z***_c_*) by maximizing the partial log-likelihood. Using the Breslow method ^43^, we maximize:

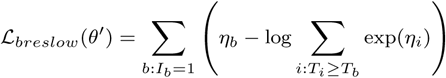

where *I_b_* = 1 indicates the occurrence of an event (death) for the patient of bulk *b*. *T_b_* represents the time of death when *I_b_* = 1, or the last contact time when *I_b_* = 0.

Using the Efron method ^44^, we maximize:

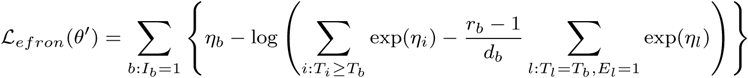

where *d_b_*is the total number of patients experiencing events simultaneously at *T_b_*, and *r_b_* is the 1-based index indicating the position of bulk *b* in tied data, with *r_b_* = 1 when *d_b_* = 1.

Using the relation 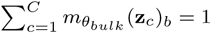, we obtain:

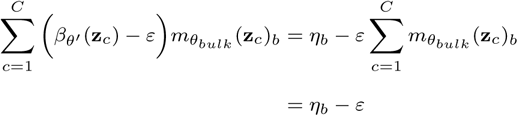

From this fact, when *η* is translated to *η* − *ɛ* in the equation *ℒ_breslow_*, we get that the loss function is invariant as following.

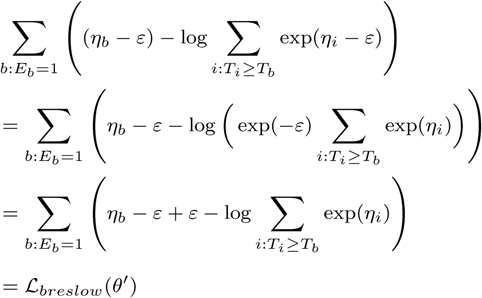

As a same manner, *ℒ_efron_*is invariant as following.

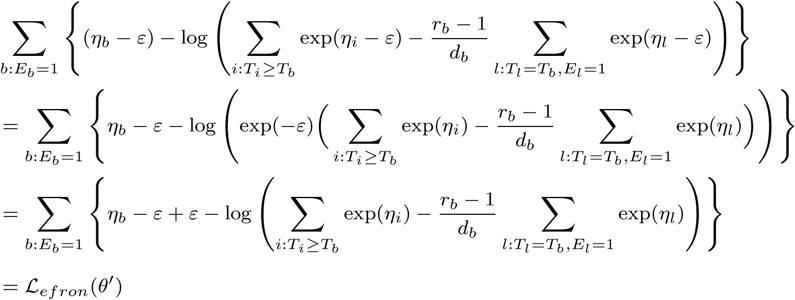

Therefore, we used *β_θ′_* (*z_c_*) only for relative comparisons between cells. In downstream analyses, we transformed *β_θ′_* (*z_c_*) by setting *ε* = min(*β_θ′_* (*z_c_*)) to ensure that the minimum value equals zero.

### Model training

The VAE was trained on 85% of the scRNA-seq data, with 10% allocated for validation and 5% for testing. The training was conducted using mini-batches of 1,000 cells, and the process was terminated when the validation loss averaged over 10 epochs failed to improve for 10 consecutive epochs. The model was optimized using AdamW with a learning rate of 0.01.

After training the VAE and fixing its parameters, the deconvolution model was trained using the same data splitting and training procedures. When the deconvolution model converged, the parameters were fixed.

For the contribution estimation, the validation and test datasets of the cells were the same as in the previous steps, but the mini-batch size for the training data was set to 500 cells. Bulk RNA-seq data were randomly split into 60% training, 20% validation, and 20% test sets to ensure that the proportion of patients with events was approximately balanced across these subsets. The training was terminated when the c-index, computed from the hazard estimates on both the cell and patient validation sets and averaged over 10 epochs, showed no improvement over 10 consecutive epochs. For this step, the AdamW optimizer was used, with a learning rate of 0.0001.

### Simulation of deconvolution and survival prediction

Pseudo-bulk samples were created using the breast cancer scRNA-seq data from Wu et al. ^45^ We first selected scRNA-seq data from the patient with the highest cell count and randomly split it in half. We used one half to create pseudobulk samples and the other half as a reference. A total of 300 pseudo-bulk samples were generated in this process. We utilized the pre-assigned ‘cell-type_minor’ annotation as cluster labels. The selection probability for each cluster was generated from a Dirichlet distribution with *α* = 1 to ensure uniform selection across cluster labels. The number of cells extracted from each cluster was generated from a multinomial distribution by using these selection probabilities. The total cell count for each pseudobulk was sampled from a discrete uniform distribution *U* {1000, 10000}.

Following Austin’s methodology ^46^, we simulated the survival time *T_b_*for each bulk sample:

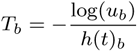

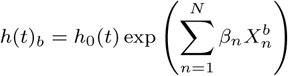

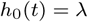

where *u_b_* ∼ *U* (0, 1) (standard uniform distribution), *λ* = 0.001, *N* is the number of ‘celltype_minor’ clusters, *β* ∼ N (0, 5*I*), and *X* represents the cell count of each cluster in each bulk sample. To evaluate the accuracy of regression coefficient estimation, we generated 100 datasets using the same 300 pseudo-bulk samples, each with different survival time settings obtained by varying the *β* values. All samples were set to be uncensored.

### scRNA-seq data preprocessing

All transcriptome count data were processed using the Scanpy Python package. For cancer reference scRNA-seq data exceeding 10,000 cells, we randomly sampled 10,000 cells. For COVID-19 reference scRNA-seq data ^32^, cells annotated as ‘COVID_status: COVID-19’ and ‘Group: Adult’ were included to ensure consistency with the bulk RNA-seq data. From these filtered cells, 10,000 cells were randomly sampled. Genes expressed in less than 1% of all cells were excluded. This gene filtering criterion was similarly applied to the bulk and spatial transcriptome datasets. We retained genes common to both scRNA-seq and bulk RNA-seq datasets. We selected the top 5,000 highly variable genes in bulk RNA-seq using scanpy.pp.highly_variable_genes and used them as inputs for scSurv.

### TCGA bulk data preprocessing

Clinical information and bulk RNA-seq data were obtained from the Genomic Data Commons (GDC) Data Portal. For patients with multiple bulk RNA-seq samples, we prioritized biopsies from primary tumors, followed by metastatic tumors if the primary tumors were unavailable. When two biopsies were obtained from the same site, the biopsy with the highest total count was selected. Samples lacking both death time and the last contact time were excluded. Following Liu et al. ^47^, we obtained sample procurement time from the ‘days_to_sample_procurement’ field. For ovarian cancer data, for which most sample procurement times were missing, we set the procurement time equal to the diagnosis time. For other cancers, we excluded data with missing sample procurement dates and calculated the survival time from the procurement time as time zero.

### COVID-19 PBMC bulk data preprocessing

Clinical information and bulk RNA-seq data were obtained from the IMPACC cohort. ^31^ This cohort includes 1164 hospitalized COVID-19 patients, aiming to investigate immune responses, disease progression, and potential biomarkers. We used bulk RNA-seq data from PBMCs. For patients with multiple samples, we selected samples collected when the patient’s respiratory status was the most severe. We excluded samples that were collected after discharge or death, along with those missing survival or discharge data. After applying these exclusion criteria, 379 patients with available PBMC bulk RNA-seq data were used as input for scSurv. In this study, we focused on survival and discharge as the outcomes because they are common endpoints in time-to-event data analysis and are relevant for understanding disease progression and recovery in hospitalized patients.

### Generalization Performance of scSurv Across 12 TCGA Cancers

We applied scSurv to 12 cancer types with over 300 patients and non-rare death events. We performed 100 different training-validation-test splits using different random seeds, applying scSurv once for each seed. To quantitatively evaluate prediction accuracy, we calculated the c-index for the training, validation, and test data using the lifelines.utils.concordance_index function from the lifelines library. For comparison, we performed a regression on the hazard function using dimensionally reduced bulk RNA-seq counts. Regression was performed using the CoxPHFitter function from the lifelines library. We set penalizer=0.01 to prevent convergence errors. Using the same training data as scSurv, we performed a hazard function regression and calculated the c-indices of the test data.

### Spatial transcriptome preprocessing

We obtained Visium data for kidney cancer from Li et al.’s ^23^ public dataset. Following their criteria, we excluded Visium spots with counts below 2,000 or above 35,000, spots with fewer than 500 genes, and spots with mitochondrial gene percentages exceeding 20%. In addition, we removed two low-quality slides (6800STDY12499504 and 6800STDY12499505) with < 500 spots. We retained the genes that were commonly detected across scRNA-seq, bulk RNA-seq, and spatial transcriptome data. We selected the top 1000 highly variable genes in bulk RNA-seq using scanpy.pp.highly_variable_genes and used them as inputs for scSurv. The cell proportions for each spot were estimated using the same method that was used for bulk deconvolution in scSurv. Using the estimated survival time contributions *β_θ′_* (**z***_c_*) and cell proportions in Visium spots *m_θ^sPot^_* (**z***_c_*), we defined the hazard function for each spot as:

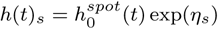

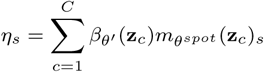

where *s* denotes the spot, *t* is time, and 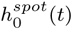 is the baseline hazard function. For visualization, we divided each spot’s hazard by the mean hazard across spots to eliminate the baseline hazard function.

### Permutation test

For permutation tests, we randomly reassigned contributions *β_θ′_* (**z***_c_*) to the hazard function among cells within each cell type cluster. We then recalculated c-indices using all patients to determine how the permutation of each cell type affected the c-index. Cell types contributing to large c-index changes were considered necessary for prognosis prediction.

### Gene set enrichment analysis

We performed gene set enrichment analysis using GSEApy ^48^. We used the ‘Reactome_2022’ database for the gene set. In the analyses of melanoma, renal cell carcinoma, and COVID-19, we used the input genes for scSurv as the background. For the pan-cancer analysis, the overlapping genes from scSurv inputs across the six cancers were used as the background.

### Cell-cell communication analysis

Following the methodology of Kojima et al. ^13^, we extracted the activated ligand-receptor relationships between identified T cells and non-T cells in the kidney cancer dataset. Using spatial transcriptome data, we first extracted cell pairs that co-localized in the same spots at rates twice as high as expected from random distribution. We then extracted pairs consisting of ‘CD8+T_cycling-G2/M’ cluster cells and other cell types from the co-localized pairs. Using genes expressed in these pairs showing expression in the top 5% percentile, we investigated activated cell-cell communication using NicheNet. ^49^

### Multiple cancer analysis

We analyzed six cancers showing c-indices above 0.5 in test data: melanoma, lung adenocarcinoma, renal cell carcinoma, hepato-cellular carcinoma, head and neck cancer, and bladder cancer. Cell type annotations were available from the original papers for all cancers except bladder cancer, which we annotated using the marker genes from Li et al ^50^. In hepatocellular carcinoma, we subclustered the ‘T/NK’ cluster into T cell and NK cell clusters using the NCR1 gene.

## CODE AND DATA AVAILABILITY

The implementation of scSurv is available at https://github.com/3254c/scSurv. The single-cell RNA sequencing datasets used in this study are available from the Gene Expression Omnibus (GEO) under accession numbers GSE129845 (kidney cancer) ^23^, GSE115978 (melanoma) ^51^, GSE129845 (bladder cancer) ^50^, GSE131907 (lung adenocarcinoma) ^52^, GSE176078 (breast cancer) ^45^, GSE132465 and GSE144735 (colorectal cancer) ^53^, GSE164690 (head and neck cancer) ^54^, GSE149614 (hepatocellular carcinoma) ^55^, and GSE183904 (gastric cancer) ^56^; from ArrayExpress under accession numbers E-MTAB-11948 (cervical cancer) ^57^ and E-MTAB-8107 (lung squamous cell carcinoma) ^58^; from Cell x Gene Explorer (Collection ID: 4796c91c-9d8f-4692-be43-347b1727f9d8) for ovarian cancer ^59^; and from COVID-19 Cell Atlas (https://covid19cellatlas.org) for COVID-19 PBMCs. Bulk RNA sequencing data and clinical information from TCGA are available from the Genomic Data Commons (GDC) Data Portal (https://portal.gdc.cancer.gov/). The bulk RNA-seq and clinical outcome data from the IMPACC cohort used in this study are accessible via the ImmPort database (https://www.immport.org) with a data access request.

## ACKNOWLEDGMENTS

We gratefully acknowledge the IMPACC study group ^31^ for providing bulk RNA-seq data and associated clinical outcome data used in this study.

## AUTHOR CONTRIBUTIONS

Y.K. conceived the method. C.M. designed and implemented the source code, performed validation studies, and conducted analyses on real-world datasets under the supervision of Y.K. and T.S. S.H. and K.A. contributed to the theoretical development of this method. D.K. contributed to the infectious disease analyses. All authors reviewed and approved the final manuscript.

---

**Figure S1:**
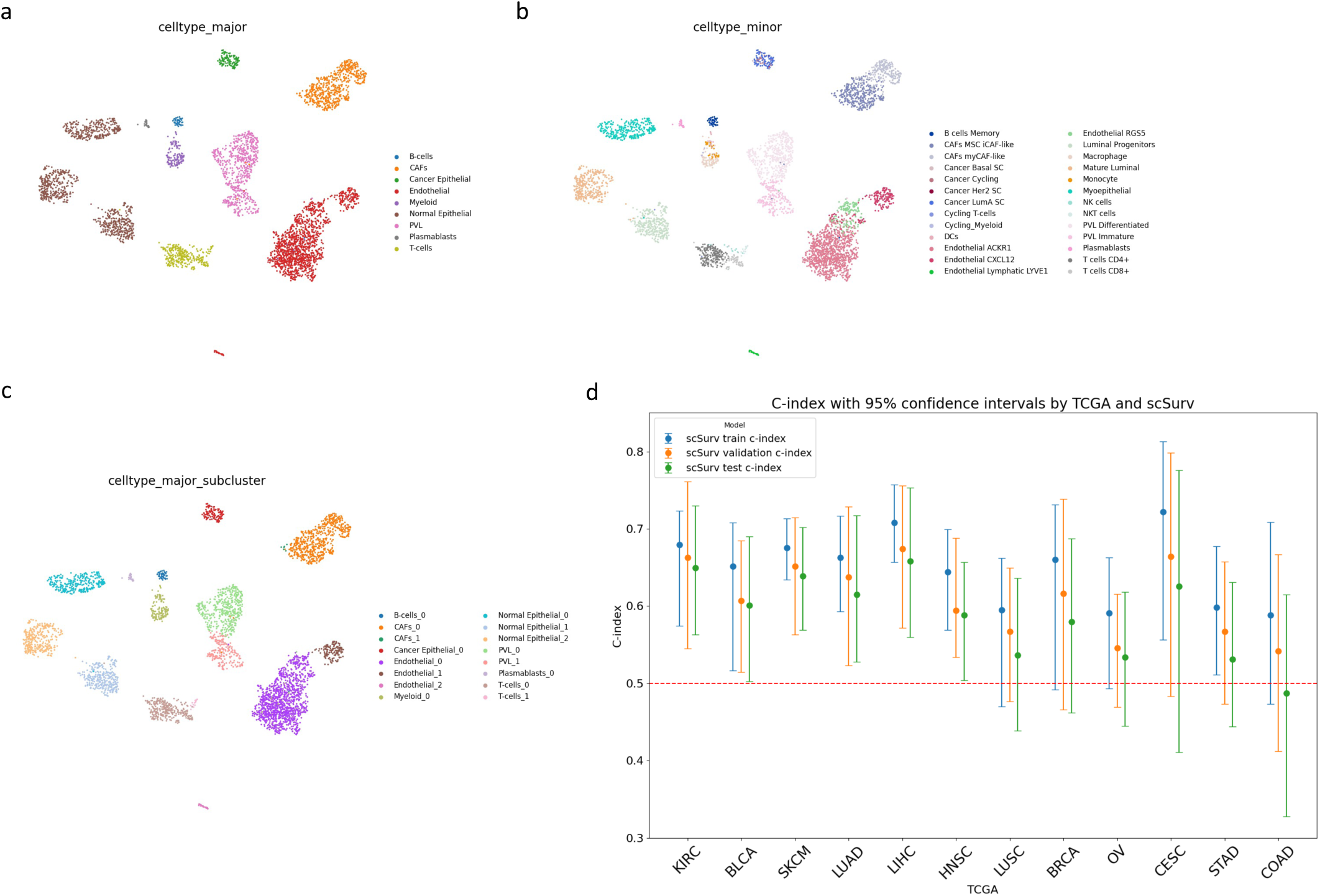
Schematic illustrations of apparatus design. (a) UMAP visualization of ‘celltype_major’ clusters. (b) UMAP visualization of ‘celltype_minor’ clusters used for generating pseudo-bulk samples in simulation. (c) UMAP visualization of ‘celltype_major’ subclusters, generated using sc.tl.leiden() with resolution=0.1. These cluster labels were used as alternative annotations for BayesPrism, CIBERSORTx, and MuSiC in simulation. (d) scSurv performance is measured by c-index across training, validation, and testing patients for 12 cancer types.

**Figure S2:**
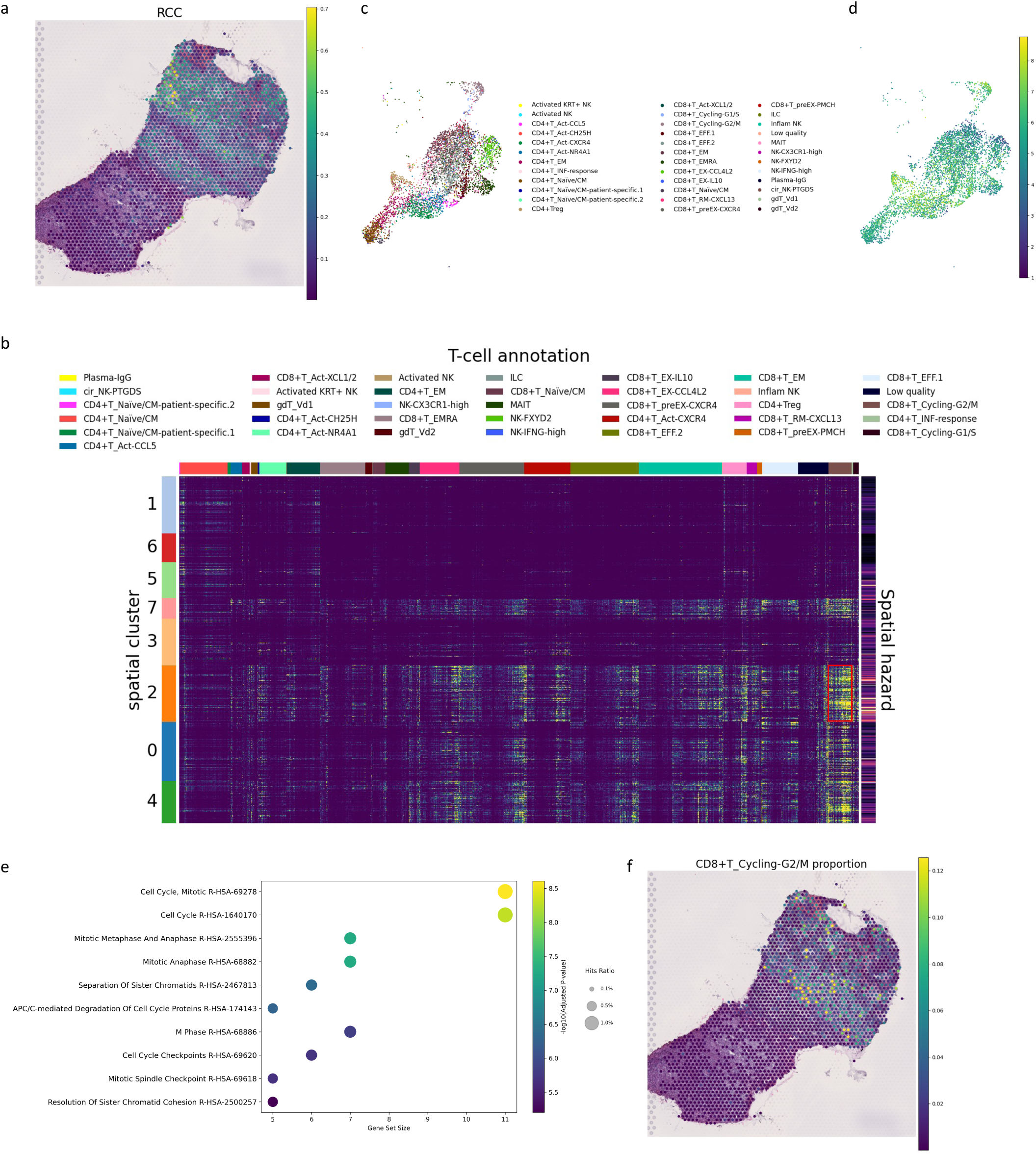
Schematic illustrations of apparatus design. (a) The proportion of cancer cells in each spot estimated by scSurv. (b) Heatmap of single-cell contributions adjusted for cell proportions in T cells. ‘CD8+T_cycling-G2/M’ cluster is specific to cluster 2. (c) UMAP visualization of T cell subtypes. (d) UMAP visualization of contributions in T cells. (e) Gene set enrichment analysis results for differentially expressed genes in the ‘CD8+T_cycling-G2/M’ cluster. (f) Spatial distribution of ‘CD8+T_cycling-G2/M’ cluster proportion.

